# Local-community network automata modelling based on length-three-paths for prediction of complex network structures in protein interactomes, food webs and more

**DOI:** 10.1101/346916

**Authors:** Alessandro Muscoloni, Ilyes Abdelhamid, Carlo Vittorio Cannistraci

## Abstract

From nests to nets intricate wiring diagrams surround the birth and the death of life. Here we show that the same rule of complex network self-organization is valid across different physical scales and allows to predict protein interactions, food web trophic relations and world trade network transitions. This rule, which we named CH2-L3, is a network automaton that is based on paths of length-three and that maximizes internal links in local communities and minimizes external ones, according to a mechanistic model essentially driven by topological neighbourhood information.

## Introduction

Recently, a pioneering study of Kovács et al. [1] empirically disclosed a link prediction principle based on paths of length three (L3) for network-based prediction of protein interactions [1] that astonishingly overcomes any rival method based on paths of length two (L2). The L3 principle was motivated by both 3D-structural and evolutionary arguments associated with protein interactions, and the associated mechanistic model (mathematical formula) introduced by Kovács et al. [1] for modelling the L3 principle in unweighted and undirected networks is:

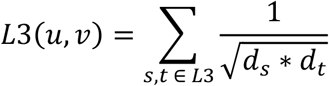

where: *u* and *v* are the two seed nodes of the candidate interaction; *s* and *t* are the two intermediate nodes on the considered path of length three; *d*_*s*_ and *d*_*t*_ are the respective node degrees; and the summation is executed over all the paths of length three. Kovács et al. motivate the penalization for the degree of *s* and *t* as follows [1]: ≪ We expect that node pairs connected by the highest number of L3 paths are most likely to be directly connected. However, high degree nodes (hubs) might induce multiple, unspecific shortcuts in the network, biasing the results. To cancel potential degree biases caused by intermediate hubs in the paths, we assign a degree-normalized L3 score to each node pair, *u* and *v* ≫.

However, although this new graceful link prediction strategy was introduced according to biologically sound rationales and was convincingly supported by empirical evidence, its mathematical definition as a mechanistic model was empirically introduced more as a matter of intuition than as derivation from current theoretical knowledge, and it actually misses any connection to already known link prediction generalized principles. *De facto*, this represents a significant conceptual limitation because it obstacles the theoretical understanding of the universal mechanisms and modelling formalisms that are behind link prediction. The importance of the L3 model discovery remains valid, but the lack of understanding has a clear negative impact on both the definition of a comprehensive theory of link prediction and the improvement/extension of L3 modelling to any type of complex network.

## Results

### Principles and modelling

#### Resource allocation on paths of length n

The first objective of this study is to understand the generalized mechanistic principle of complex network self-organization behind the mathematical formula introduced by Kovács et al. [1] for modelling the L3 principle in unweighted and undirected networks. And, what is the first important discovery?

The first important finding of this study is surprisingly banal: the generalized principle behind the L3 formula proposed by Kovács et al. [1] is already well known in network science, in particular in link prediction, and lies in only two words: *resource allocation* (RA) [2]. In order to prove it we need few and simple mathematical steps. The basic formula of the RA model on a path of length two (which from here forward we will call RA-L2) is:

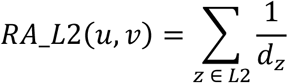

where: *u* and *v* are the two seed nodes of the candidate interaction; *z* is the intermediate node on the considered path of length two; *d*_*z*_ is the respective node degree; and the summation is executed over all the paths of length two.

In order to generalize to paths of length *n* > 2, we need an operator that merges the single contributes of each weighted (in this case degree-penalized) common neighbour on the path of length *n*. If, without lack of generality, we use as merging operator the geometrical mean (which is a choice for designing a robust estimator, since if only one common neighbour on the path has a low weight, then the whole path is penalized), we derive the following generalized formula for paths of length *n*:

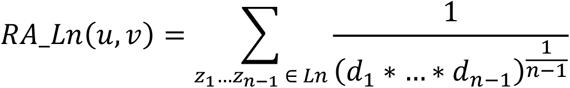

where: *u* and *v* are the two seed nodes of the candidate interaction; *z*_1_…*z*_*n*-1_ are the intermediate nodes on the considered path of length *n*; 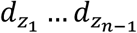 are the respective node degrees; and the summation is executed over all the paths of length *n*.

For paths of length *n* = 3, the general formula given above becomes clearly equal to L3, which indeed extends the resource allocation principle on paths of length three, therefore from here forward we will call L3 with the new name of RA-L3.

#### Local-community-paradigm and Cannistraci-Hebb automata on paths of length *n*

The second objective of this study is to extend to paths of length *n* also the local community paradigm (LCP) theory and the associated network automata model for link prediction that at the moment are defined only on L2. The motivation to generalize LCP methods to L3 is that empirical evidences provided by several studies [3]–[9] in link prediction, and confirmed also by Kovács et al. [1], show that Cannistraci resource allocation (CRA) - which is putatively the local community extension of RA-L2 - outperforms both RA-L2 and the large majority of other neighbourhood-based mechanistic and parameter-free L2-models. Hence, the question is whether also CRA-L3 would outperform RA-L3. To make so, we need before to review and extend the LCP theory to paths of length *n* and then we need to define the mathematical formula of the new CRA-L3 model that arises according to this generalization. In order to implement this plan, at first we need to recall the basic rationale behind the LCP theory.

In 1949, Donald Olding Hebb advanced a *local learning rule* in neuronal networks that can be summarized in the following: neurons that fire together wire together [10]. In practice, the Hebbian learning theory assumes that different engrams (memory traces) are memorised by the differing neurons’ cohorts that are co-activated within a given network. Yet, the concept of wiring together was not further specified, and could be interpreted in two different ways. The first interpretation is that the connectivity already present, between neurons that fire together, is reinforced; whereas the second interpretation is the emergence and formation of new connectivity between non-interacting neurons already embedded in an interacting cohort. In 2013 Cannistraci et al. [3] noticed that, considering only the network topology, the second interpretation of the Hebbian learning could be formalized as a mere problem of topological link prediction in complex networks. The rationale is the following. The network topology plays a crucial role in isolating cohorts of neurons in functional communities that naturally and preferentially can perform local processing, by virtue of this predetermined local-community topological organization. In practice, the local-community organization of the network topology creates a physical and structural ‘energy barrier’ that allows the neurons to preferentially fire together within a certain community and therefore to add links inside that community, implementing a type of local topological learning. In few words: the local-community organization influences (by increasing) the likelihood that a cohort of neurons fires together because they are confined in the same local community, consequently also the likelihood that they will create new connections inside the community is increased by the mere structure of the network topology. Inspired by this intuition, Cannistraci et al. [3] called this local topological learning theory *epitopological learning,* which stems from the second interpretation of the Hebbian leaning. The definition was not clearly given in the first article [3] that was quite immature, and therefore we now provide an elucidation of the concepts behind this theory by offering new definitions. *Epitopological learning* occurs when cohorts of neurons tend to be preferentially co-activated, because they are topologically restricted in a local community, and therefore they tend to facilitate learning by forming new connections instead of merely retuning the weights of existing connections in the local community. As a key intuition, Cannistraci et al. [3] postulated also that the identification of this form of learning in neuronal networks was only a special case, hence the *epitopological learning* and the associated *local-community-paradigm (LCP)* were proposed as local rules of learning, organization and link-growth valid in general for topological link prediction in any complex network with LCP architecture [3]. On the basis of these ideas, they proposed a new class of link predictors that demonstrated - also in following studies of other authors - to outperform many state of the art local-based link predictors [3]-[9], [11] both in brain connectomes and in other types of complex networks (such as social, biological, economical, etc.). In addition, they proposed that the local-community-paradigm is a necessary paradigm of network organization to trigger epitopological learning in any type of complex network. In conclusion, the LCP originated from the initial idea to explain how the network topology indirectly influences the process of learning a memory by adding new connections in a network of neurons, and consequently it was generalized to advocate mechanistic modelling of topological growth and self-organization in real monopartite [3] and bipartite [12] complex networks, with a significant impact also on prediction of drug-target interactions exploiting exclusively bipartite network topology [13]. A recent study of Narula et al. [14] shows that local-community-paradigm and epitopological learning can enhance our understanding of how local brain connectivity is able to process, learn and memorize chronic pain [14]. And how can this be exploited also in the domain of prediction of protein interactions?

Protein interactomes display a clear LCP architecture [3], where protein complexes are confined in local and topologically isolated network structures, which are often coincident with functional network modules that play a crucial role in molecular circuits. The key generalized idea behind the LCP network architecture is that, for instance, a local community of neurons or proteins should take functional advantage of being confined in a local assembly of operational units. Each local assembly - if it is properly activated by an external signal coming from another region of the network - performs a functional operation by means of a structural remodelling of the internal connectivity (named iLCL in Fig. 1) between the operational units that are embedded in the network local community. The systems supported by LCP network architecture are very dynamic and react to a stimulus with a local plastic remodelling. In case of operational units such as neurons, the local community remodelling can implement for instance a learning process. Instead, In case of operational units such as proteins, the local community remodelling is necessary to implement for instance a biological process, which emerges by the molecular-complex rearrangement in the 3D space.

**Figure 1.**
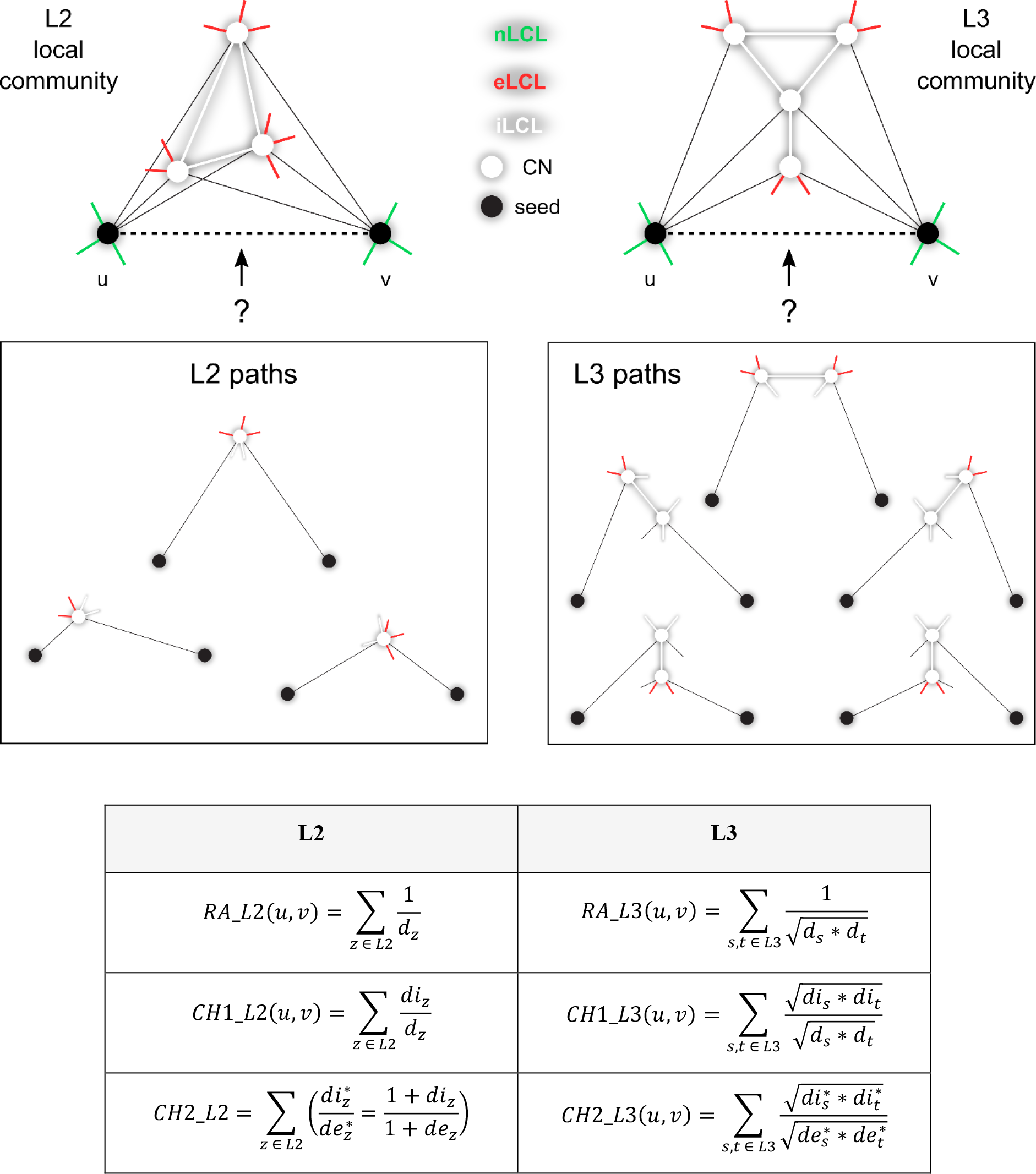
Cannistraci-Hebb epitopological rationale. The figure shows an explanatory example for the topological link prediction performed using the L2 or L3 Cannistraci-Hebb epitopological rationale. The two black nodes represent the seed nodes whose non-observed interaction should be scored with a likelihood. The white nodes are the L2 or L3 common-neighbours (CNs) of the seed nodes, further neighbours are not shown for simplicity. The cohort of common-neighbours and the iLCL form the local-community. The different types of links are reported with different colours: non-LCL (green), external-LCL (red), internal-LCL (white). The set of L2 and L3 paths related to the given examples of local communities are shown. At the bottom, the mathematical description of the L2 and L3 methods considered in this study are reported. Notation: *u*, *v* are the seed nodes; *z* is the intermediate node in the L2 paths; *s*, *t* are the intermediate nodes in the L3 paths.

The previous conceptual and mathematical formalizations of the LCP-theory were immature and put more emphasis on the fact that the information content related with the common neighbour nodes should be complemented with the topological information emerging from the interactions between them (the iLCL in Fig. 1). However, here we would like to remark that the local isolation of the operational units in the different local communities is equally important to carve the LCP architecture in the network, and this is guaranteed by the fact that the common neighbours minimize their interactions external to the local community (the eLCL in Fig. 1). This minimization forms a sort of topological energy barrier, which in turn confines the signal processing to remain internally to the local community. Hereafter, we will revise the LCP idea and its mathematical formalization in order to explicitly take into account also the *minimization of the external* links (the eLCL in Fig. 1). However, in this article we will discuss the implications of this theoretical revision not only on modelling of protein interactomes, but also on other types of networks.

Recently, Muscoloni et al. [15] discussed how local parameter-free mechanistic models to predict link-growth in complex networks (such as the common neighbours and LCP-based indices discussed in the previous section) can be interpreted as network automata that compute the likelihood to close ‘local rings’ in the network whenever a link is missing in the topology (see Suppl. Fig. 1 for visual clarification). The *local ring* is the closure of a ‘local tunnel’ obtained by adding to the topology the missing link for which the likelihood to appear is computed. The *local tunnel* is the ensemble of all the local paths (which can be the smallest shortest-paths definable on a given network topology or the paths of a fixed arbitrary length) that connect two nonadjacent nodes (extremities of the tunnel), and the common neighbours are all the nodes embedded in the tunnel structure, therefore they represent an estimation of the size of the tunnel (see Fig. 1 for an example). The study of Muscoloni et al. [15] discusses also that the Cannistraci-resource-allocation (CRA) is a local-ring network automaton model that seems strongly related and able to predict the growth of network topology which is associated to hyperbolic geometry.

For the CRA network automaton model the likelihood of a new link to appear is function not only of the number of common neighbours, but also function of the iLCLs and eLCLs (Fig. 1). Therefore, the mathematical formula of CRA in L2 is:

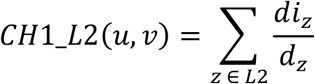

where: *u* and *v* are the two seed nodes of the candidate interaction; *z* is the intermediate node on the considered path of length two; *d*_*z*_ is the respective node degree; *di*_*z*_ is the respective internal node degree (number of iLCL); and the summation is executed over all the paths of length two.

However, looking at this formula, it is evident that the principle behind CRA is not a resource allocation penalization at all. On the contrary, the model is based on the common neighbours’ rewards of the internal links (iLCL) balanced by the penalization of the external links only. Hence, the current name is misleading. Since this model is a generalization and reinterpretation of a Hebbian learning local-rule to create new topology in networks, we decide to rename CRA as L2-based Cannistraci-Hebb network automaton model number one (CH1-L2), rather than Cannistraci-Resource-Allocation (CRA), as it was named in the previous articles [3]–[9], [11]–[13]. Yet, we have to admit that the formula of CH1-L2 is conceptually and mathematically awkward. If we want to design a model that is based on the minimization of the eLCL and the maximization of the iLCL, why not to write it directly and explicitly? This is what we propose to realize with the definition of a new model that we name CH2-L2 and that theoretically should be a straightforward mathematical formalization to express the principle or rule of topological self-organization behind the Cannistraci-Hebb network automaton modelling. The formula of CH2-L2 is:

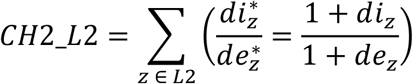

where: *u* and *v* are the two seed nodes of the candidate interaction; *z* is the intermediate node on the considered path of length two; *di*_*z*_ is the respective internal node degree (number of iLCL); *de*_*z*_ is the respective external node degree (number of eLCL); and the summation is executed over all the paths of length two. Note that a unitary term is added to the numerator and denominator to avoid the saturation of the value in case of iLCL or eLCL equal to zero.

Suppl. Fig. 1 suggests a geometrical interpretation about how the CH network automata models work in a monopartite topology. The local-tunnel (which is formed in a hidden highdimensional geometrical space that here we project in the hyperbolic disk for simplifying the visualization, provides a route of connectivity between the two nonadjacent nodes. Although in the hyperbolic disk visualization the local-tunnel has a shape that resembles the hyperbolic distances (curved toward the centre), we clarify that the proposed term ‘tunnel’ is only an idealized definition; in fact the shape of this bridging structure could also geometrically look as a chamber or basin, but this does not change the meaning of the definition. The higher the number of common neighbours, the higher the size of the local-tunnel. For each common neighbour, the higher the number of iLCLs in comparison to the eLCLs, the more the shape of the local-tunnel is well-defined and therefore its existence confirmed. Therefore, in link prediction, CH models estimate a likelihood that is proportional both to the size of the local-tunnel and to the extent to which the local-tunnel exists. On the other hand, the common neighbour network automaton model only estimates a likelihood proportional to the size of the local-tunnel, which is a significant limitation. Furthermore, the local-community represents the central chunk of the local-tunnel exclusively formed by CNs and iLCLs. The local-community represents a bridging structure which is fundamental to transfer the information between the two seed nodes.

In order to formulate CH1 and CH2 in paths of length *n*, we need to define what is the local-community in paths of length *n*. To this aim, it is easier to provide an example for paths of length three as represented in the Fig. 1. The common neighbours are all the intermediate nodes touched by any path of length three that connect the two seed nodes *u* and *v*. The iLCL are the links between common neighbours that are attached to different seed nodes, to guarantee a direct flux of information that forms a local-tunnel between the seed nodes (Fig 1). On the contrary, an example of links that should not be considered as iLCL although they connect common neighbours are provided in Suppl. Fig. 2. These links are not iLCL because they do not participate directly to deliver flux of information from one seed node to the other one, in fact they create a bridge between two nodes that are exclusively neighbours of the same seed node. The local-community is the structure that is posed at the centre of the tunnel formed by the L3 paths and it is composed of the common neighbours and the iLCL. Hence, the mathematical formulae for the CH1 and the CH2 models for paths of length *n* are:

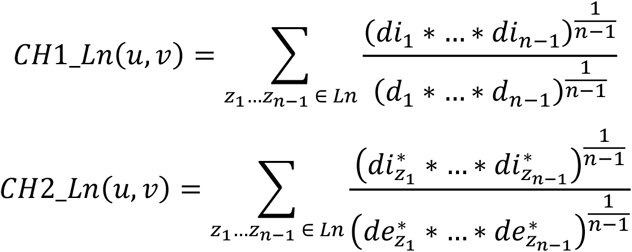

where: *u* and *v* are the two seed nodes of the candidate interaction; *z*_1_…*z*_*n*-1_ are the intermediate nodes on the considered path of length *n*; 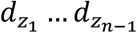 are the respective node degrees; 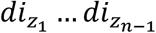 are the respective internal node degrees (number of iLCL); 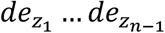 are the respective external node degrees (number of eLCL); terms with an asterisk as superscript indicate that a unitary value is added 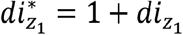 and 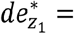 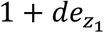; and the summation is executed over all the paths of length *n*.

The respective formulae of CH1-L3 and CH2-L3 are provided in Fig. 1.

At the end of this paragraph we want to stress that till now we never spoke about the triadic closure principle because we believe that common neighbours are not associated to any triadic closure and that this is in our opinion a misleading principle. In our opinion, a correct principle to define common neighbours is in relation to the definition of local paths of length *n*. The common neighbours are in general, according to our proposed definition, all the intermediate nodes touched by any path of length *n* between two nodes in the network. In case of paths of length 2 this is specifically coincident with triadic closure, but in case of paths of length *n* we can generally define the operation of ‘local ring’ closure.

### Experimental evidences

#### Protein-protein interaction networks

Our initial experiments are based on 10% random removal and re-prediction of the original protein network topology, because as we show in Suppl. Fig. 3-5 the removal of 50% of the original links brings to a significant aberration of the network features.

The first important finding of this study is that the CH2-L3 model performs significantly better than RA-L3 (Fig. 2), and similarly CH2-L2 performs better than RA-L2 (Suppl. Fig. 6) when we consider link prediction of protein interactions. This implies that in general (regardless of L3 or L2 implementation of the model) the CH reward/penalization strategy is a principle of selforganization that works better than the RA penalization, which was the original principle behind the L3 model proposed by Kovács et al. [1]. In addition, CH2-L3 performs better than CH1-L3 (Suppl. Tables 2-3), and this result indicates that our intuition to design CH2-L3, which is a more straightforward mathematical formula for the CH principle, was well-posed. Astonishingly, the fact that in general L3 are better than L2 approaches is not confirmed by gene ontology (GO) evaluation (Suppl. Tables 4-9), where CH2-L2 and CH1-L2 seem the best methods for prediction of missing links evaluated by their relevance according to GO annotations. Unfortunately, network topology prediction evaluation and GO evaluation are in disagreement on what is the best strategy. This implies that further tests are required to investigate the reason of this disagreement. Kovács et al. [1] speculated that the legitimacy of using GO terms or functional annotations to evaluate the quality of the predicted physical interactions should be questioned. In order to face this contradictory finding, we considered the human (*H. sapiens*) protein network released by Menche et al. [16], which comprises 13460 proteins and 138427 physical interactions experimentally documented in human cells, including protein-protein and regulatory interactions, metabolic pathway and kinase-substrate interactions, representing one of the largest and completed blueprints of the human protein interactome [16]. But, also this further test provides contradictory results. On this valuable network, we actually find that the best method for prediction of protein interactions according to 10% random removal and re-prediction is CH2-L2, which significantly outperforms all the other methods, whereas according to the GO evaluation the best method is CH2-L3 (Suppl. Fig. 7). This result is crucial. It might suggest that either the network of Menche et al. [16] has some issues in the way it was assembled or the other networks are incomplete and noised. A careful analysis reveals that the average degree and average clustering of c network are respectively 20.6 and 0.17, which are significantly higher than the respective values for all the other networks that are on average 6.3 and 0.07. The natural follow up is to build synthetic networks on which we could change the average degree, average clustering and replicate the contradictory results obtained on real networks. Therefore, we performed also link prediction on synthetic networks generated with the nonuniform popularity-similarity-optimization model (nPSO) [17], [18], which is a recently introduced random geometrical graph generative model that allows to build networks in the hyperbolic space with clustering, small-worldness, scale-freeness, rich-clubness and a tailored community structure. We fixed the size to 1000 nodes and the exponent of the power-law degree distribution to 3. We tuned the average degree, the average clustering and community number. The important discovery of evaluating the 10% removal and re-prediction on synthetic networks is that L2 methods (with CH2-L2 confirming to be the best) perform better in general for high average clustering values regardless of variations on the other parameters. This makes sense because high clustering means more triangles (higher tendency to L2 behaviour) between nodes that are close in the geometrical space. Whereas L3 methods (with CH2-L3 that confirms to be the best) are better for networks that present simultaneously low average degree and very low average clustering, which indicates that the hyperbolic geometry is heavily compromised by randomness due to high temperature of the Fermi-Dirac connectivity probability distribution in the nPSO model (Fig. 3, left panels). This result might indicate that when networks with a hidden hyperbolic geometry and low average degree are noised by a consistent amount of random interactions (which represent false positives and cause a significant reduction of the average clustering), then L3 methods outperform L2 ones, which unfortunately is the exact scenario we observed for the majority of protein interactomes considered in this study. On the other hand, in networks with high average degree (around 28) and low clustering coefficient (around 0.17), it is clear from the plots in Fig. 3 (right panels) that L2 methods are better than L3, and this result is in accordance with what we found on the network of Menche et al. [16]. Hence, with these experiments on synthetic networks we could reproduce and justify the results obtained (by 10% removal and re-prediction evaluation) for both the Menche et al. [16] and the other protein interactomes. The open issue, which we leave for investigation in future studies, is to understand why GO-evaluation offers a result that is conceptually opposite to the one obtained by 10% removal and re-prediction evaluation. In particular, why on protein interactomes with low average degree, GO-evaluation points out CH2-L2 as the best method for prediction of new interactions and on the contrary on the Menche et al. [16] network, which has a significantly higher average degree than the previous ones, GO-evaluation suggests that CH2-L3 is the best method? Altogether, we can conclude that L3 methods seem to provide a promising principle to overcome the conceptual limitations of L2 ones on protein interactomes. In the majority of the cases we confirm the impressive performance of L3 methods reported by Kovács et al. [1] for topological prediction of protein interactions in tests based on 10% random removal and re-prediction of links. However, we also support with experimental evidences an important new finding that is the improvement provided by Cannistraci-Hebb network automata modelling (in particular the model CH2-L3), that significantly outperforms the previous RA-L3 modelling provided by Kovács et al. [1].

**Figure 2.**
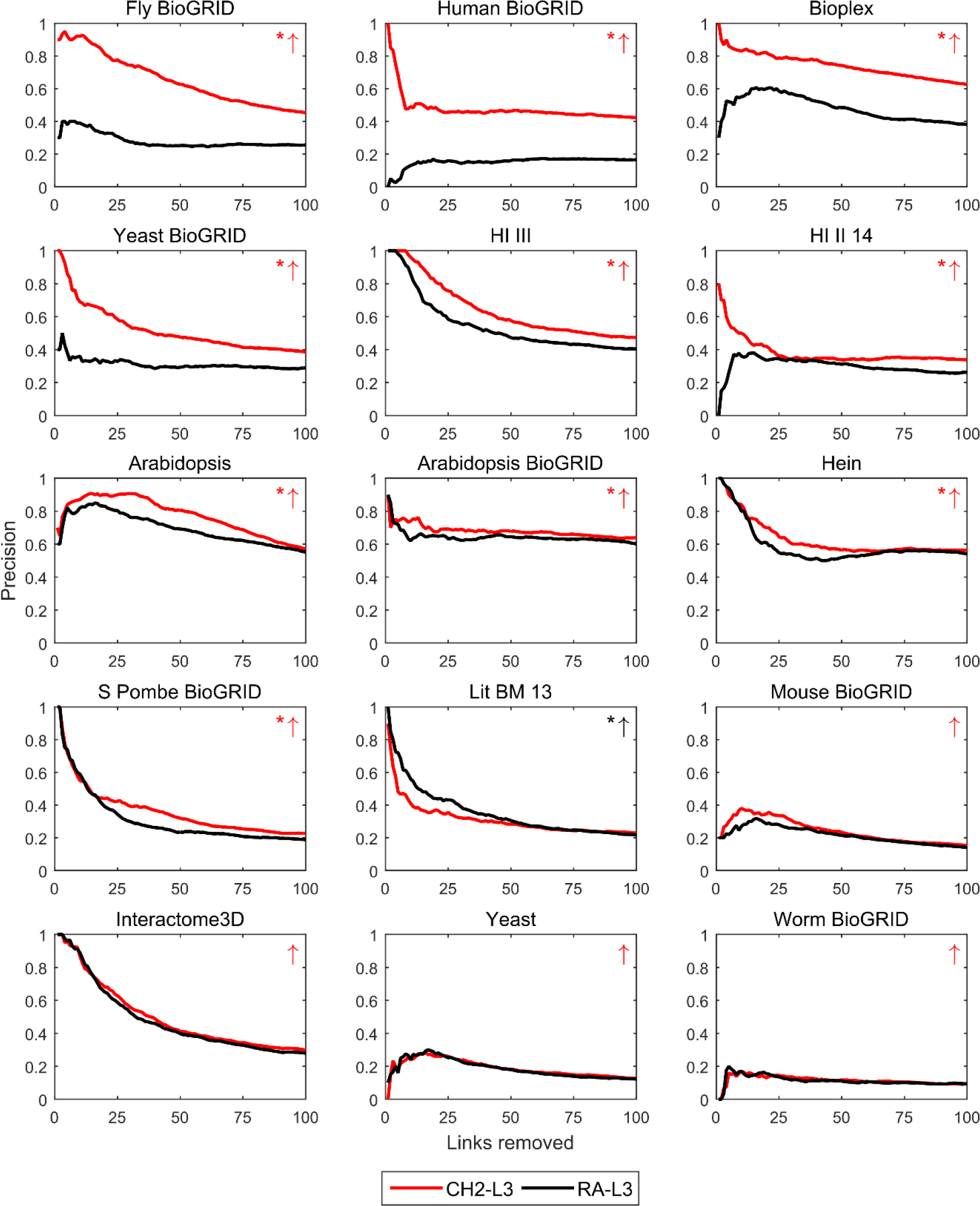
Precision curve evaluation for PPI networks: L3-based methods. For each PPI network, 10% of links have been randomly removed (10 repetitions) and the algorithms have been executed in order to assign likelihood scores to the non-observed links in these reduced networks. In order to evaluate the performance, the links are ranked by likelihood scores and the precision is computed as the percentage of removed links among the top-*r* in the ranking, for each *r* from 1 up to 100 at steps of 1. The plots report for each network the precision curve (averaged over the 10 repetitions) for the L3-based link prediction methods RA-L3 and CH2-L3. For each network, a red or black arrow is reported on the top-right of the subplot in order to indicate respectively if the average area under precision curve (AUP) is higher or lower for CH2-L3 with respect to RA-L3. A permutation test for the mean AUP has been computed, and an asterisk is reported next to the arrow in case of statistical difference (p-value ≤ 0.05). Error bars are not shown because negligible. The networks are ordered (top-down, left-right) by increasing p-value and, in case of tie, by decreasing absolute difference of mean AUP between the two methods.

**Figure 3.**
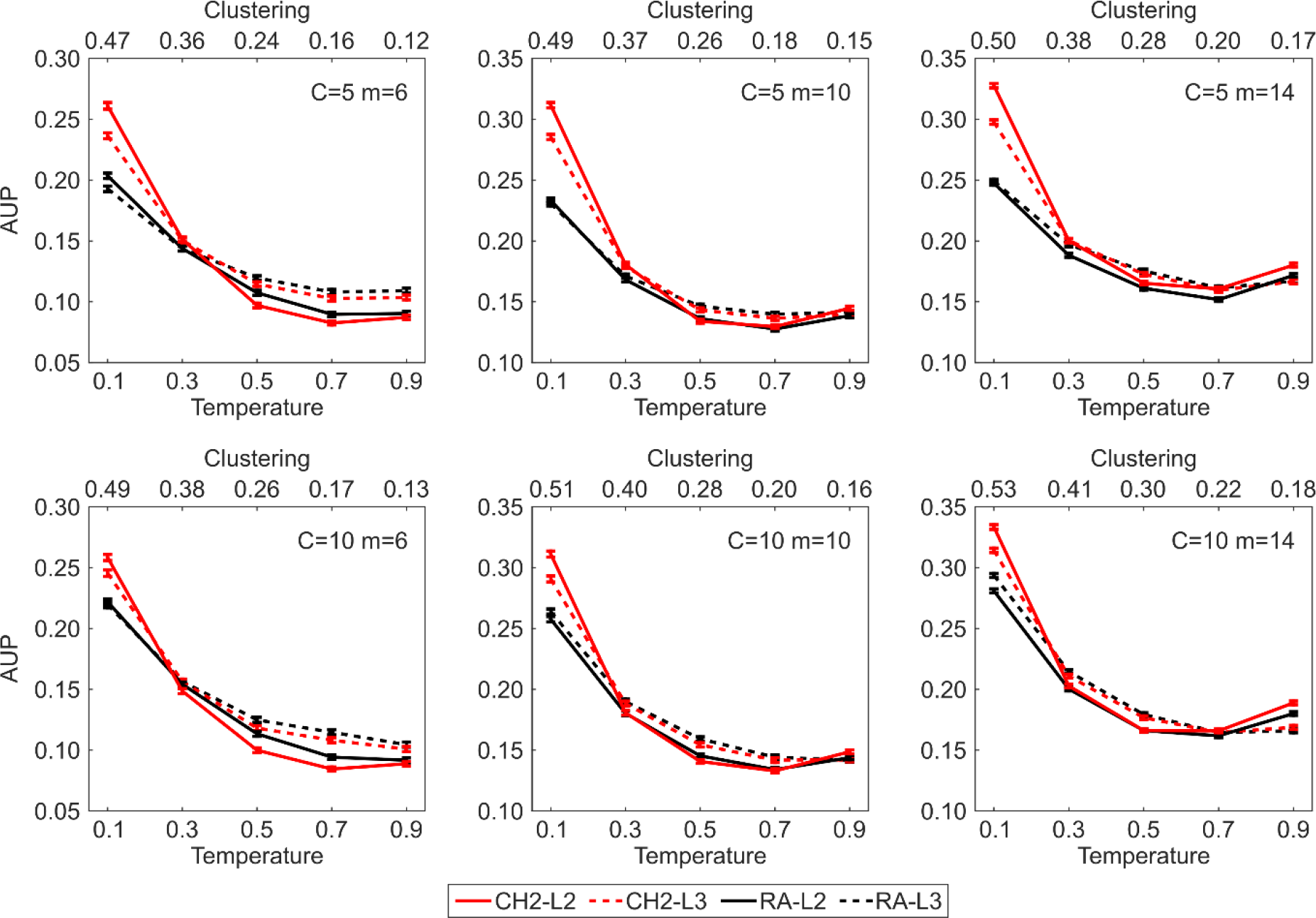
AUP evaluation for nPSO networks. Synthetic networks have been generated using the nPSO model with parameters *N* = 1000, *m* = [6, 10, 14], *T* = [0.1, 0.3, 0.5, 0.7, 0.9], *Y* = 3 and angular coordinates sampled according to a Gaussian mixture distribution with equal proportions and components *C* = [5, 10]. For each combination of parameters, 100 networks have been generated. For each network 10% of links have been randomly removed and the algorithms have been executed in order to assign likelihood scores to the non-observed links in these reduced networks. In order to evaluate the performance, the links are ranked by likelihood scores and the area under precision curve (AUP) is computed for the top-*r* removed links in the ranking, where *r* is the total number of links removed. The plots report for each parameter combination the mean AUP and standard error over the random repetitions.

#### Food webs

The second important finding of this study is that food webs seem a network class on which the result provided by L3 methods is indisputably the best (Suppl. Table 10), with CH2-L3 that again significantly outperforms RA-L3 also on the 10% removal and re-prediction evaluations performed on this networks (Fig. 4). The motivation for this new finding in respect to the study of Kovács et al. [1] that was restricted to protein interactomes is provided hereafter. In food webs two species that are at the same trophic level have a low likelihood to be predator of each other, regardless of the fact that they share many predators and/or preys. This means that the common neighbours, and therefore a principle based on L2 trophic pattern, is not the prevalent one on food webs. Instead, two species that are at different trophic levels are more likely to interact, in particular when the trophic levels are adjacent. Since L3 paths tend to have the seed nodes in different trophic levels (Suppl. Fig. 9), a principle based on L3 trophic pattern is much more valid for food webs. For further details on this subject please refer to the detailed explanation provided in Suppl. Information.

**Figure 4.**
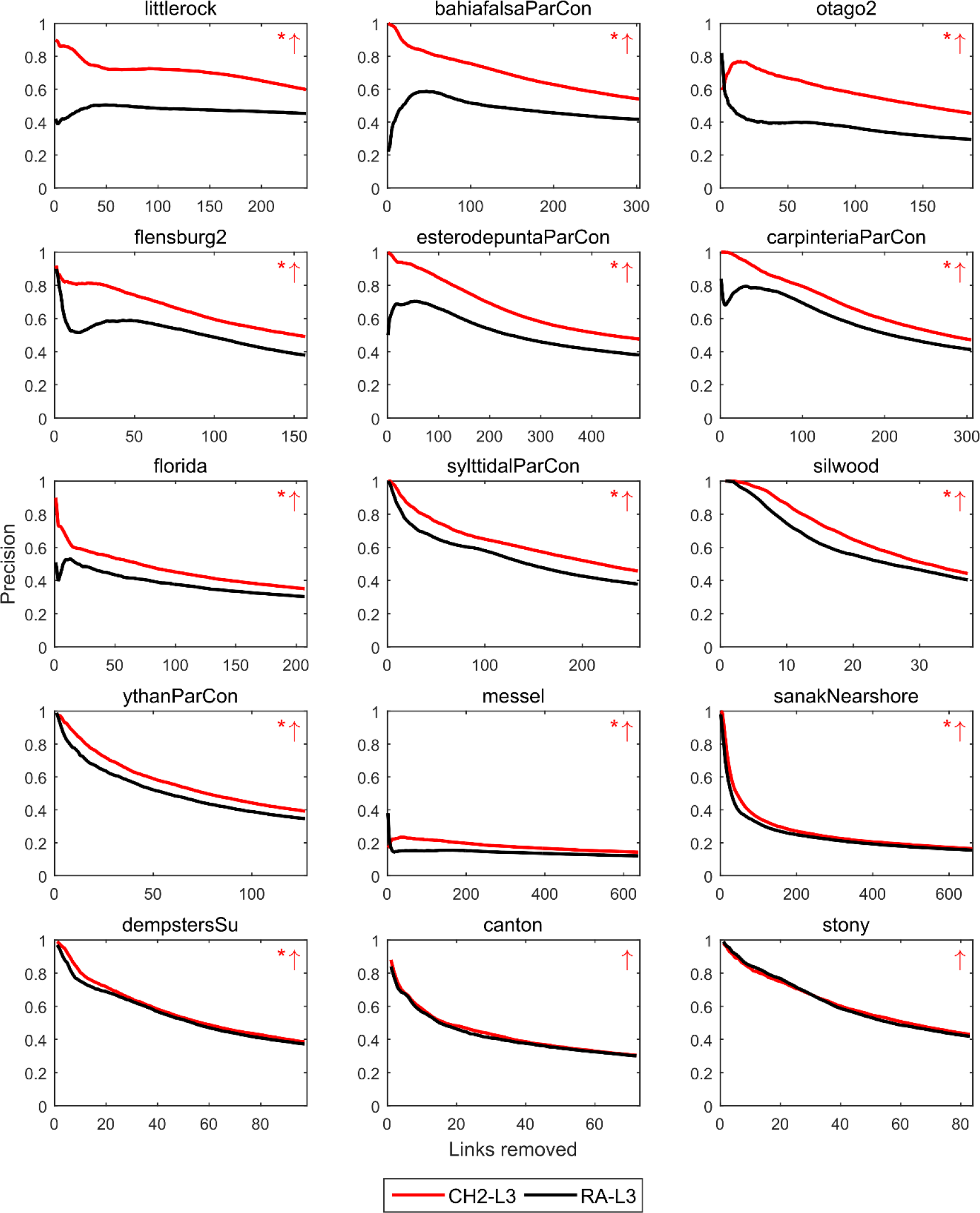
Precision curve evaluation for food webs: L3-based methods. For each food web, 10% of links have been randomly removed (100 repetitions) and the algorithms have been executed in order to assign likelihood scores to the non-observed links in these reduced networks. In order to evaluate the performance, the links are ranked by likelihood scores and the precision is computed as the percentage of removed links among the top-*r* in the ranking, for each *r* from 1 up to the total number of links removed, at steps of 1. The plots report for each network the precision curve (averaged over the 100 repetitions) for the L3-based link prediction methods RA-L3 and CH2-L3. For a matter of space, only 15 selected networks are shown (the largest networks, avoiding redundancy of geographical location). For each network, a red or black arrow is reported on the top-right of the subplot in order to indicate respectively if the average area under precision curve (AUP) is higher or lower for CH2-L3 with respect to RA-L3. A permutation test for the mean AUP has been computed, and an asterisk is reported next to the arrow in case of statistical difference (p-value ≤ 0.05). Error bars are not shown because negligible. The networks are ordered (top-down, left-right) by increasing p-value and, in case of tie, by decreasing absolute difference of mean AUP between the two methods.

#### Trade networks

The third important finding of this study is that also world international trade networks - for which it is known that the common neighbours principle is not effective [19] - follow a principle of self-organization according to which L3 is offering better link prediction performance than L2 (Fig. 5 and Suppl. Table 12). Since these networks are quite dense, link prediction is generally higher for all the methods, however also in the world trade networks CH2-L3 significantly outperforms RA-L3.

**Figure 5.**
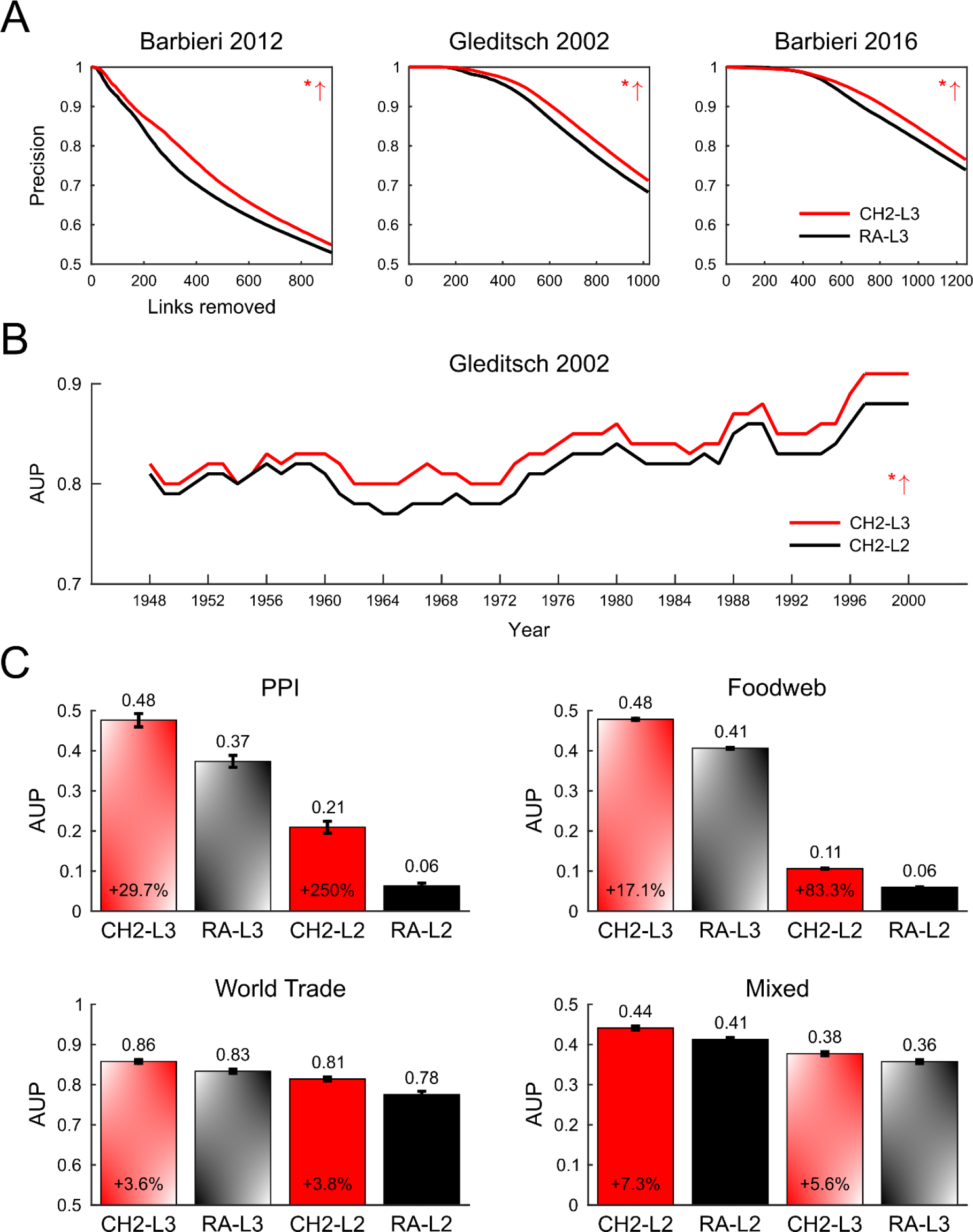
Evaluation for world trade networks and summary evaluation for real networks. (A) For each world trade network, 10% of links have been randomly removed (100 repetitions) and the algorithms have been executed in order to assign likelihood scores to the non-observed links in these reduced networks. In order to evaluate the performance, the links are ranked by likelihood scores and the precision is computed as the percentage of removed links among the top-*r* in the ranking, for each *r* from 1 up to the total number of links removed, at steps of 1. The plots report for each network the precision curve (averaged over the 100 repetitions) for the L3-based link prediction methods RA-L3 and CH2-L3. For each network, a red or black arrow is reported on the top-right of the subplot in order to indicate respectively if the average area under precision curve (AUP) is higher or lower for CH2-L3 with respect to RA-L3. A permutation test for the mean AUP has been computed, and an asterisk is reported next to the arrow in case of statistical difference (p-value ≤ 0.05). Error bars are not shown because negligible. The networks are ordered by increasing p-value and, in case of tie, by decreasing absolute difference of mean AUP between the two methods. (**B**) The same procedure described in (A) has been applied to the *Gleditsch 2002* networks available for each year from 1948 to 2000. Note that the networks in (A) are related to the last year for which information is available. The plot reports for each year the AUP (averaged over the 100 repetitions) for the link prediction methods CH2-L3 and CH2-L2. The red arrow and the asterisk have the same meaning as in (A). (**C**) Each barplot reports the mean AUP and standard error computed over all the networks and repetitions. Note that for the PPI networks the repetitions are 10 (due to the higher computational time required) and the AUP is computed for the top-100 removed links, whereas for the other networks (food webs, world trade networks and mixed real networks) the repetitions are 100 and the AUP is computed for the top-*r* removed links in the ranking, where *r* is the total number of links removed. The percentage of improvement of each CH2-method with respect to the corresponding RA-method is reported.

#### Mixed real networks

The fourth important result is that considering a plethora of mixed real networks of different nature (25 in total) the L2 principle seems the most common in comparison to L3, in fact L2 methods and in particular CH2-L2 is outperforming all the other methods in this wide evaluation analysis of mixed real networks.

## Discussion

We confirm that L3 methods seem to provide a significant improvement in respect to L2 methods for prediction of protein interactions, however network topology prediction evaluation and GO performance evaluation are in disagreement about which is the best principle between L2 and L3. This implies that further studies should investigate the reason of this disagreement. On the contrary, food webs are a class of networks on which it seems clear that L3 is a key rule of self-organization to reproduce the network topology. The same can be also confirmed for world trade networks, here the improvement is still significant but of smaller entity than on food webs, because these kinds of networks are very dense, therefore all the link predictors offer by default a good link prediction performance. Furthermore, according to our experimental evidences, the most important finding is that the local community paradigm theory and the derived Cannistraci-Hebb network automata provide a significant modelling and performance improvement on resource allocation network automata both in L2 and L3, when applied to diverse types of complex networks such as the ones investigated in this article.

## Methods

The link prediction evaluation based on 10% removal and re-prediction of the original network links is explained in the figure legend of each computational experiment. Whereas, for the gene ontology evaluation we refer to these articles that contain the details about the procedure to follow [3], [20].

## Hardware and software

MATLAB code has been used for all the simulations, carried out partly on a workstation under Windows 8.1 Pro with 512 GB of RAM and 2 Intel(R) Xenon(R) CPU E5-2687W v3 processors with 3.10 GHz, and partly in the ZIH-Cluster Taurus of the TU Dresden.

## Acknowledgements

We thank Alexander Mestiashvili and the BIOTEC System Administrators for their IT support, Gloria Marchesi for the administrative assistance and the Centre for Information Services and High Performance Computing (ZIH) of the TUD.

## Author contributions

CVC invented the principle and modelling of the article and designed the numerical experiments. AM implemented the code, recovered the data and performed the computational analysis with the help of IA. All the authors analysed and interpreted the results. CVC wrote the draft of the article and AM proposed his amendments to arrive to the final draft. CVC designed the figures and AM realized them. CVC planned, directed and supervised the study.

